# Identification of loci where DNA methylation potentially mediates genetic risk of type 1 diabetes

**DOI:** 10.1101/248260

**Authors:** Jody Ye, Tom G Richardson, Wendy L McArdle, Caroline L Relton, Kathleen M Gillespie, Matthew Suderman, Gibran Hemani

## Abstract

The risk of Type 1 Diabetes (T1D) comprises both genetic and environmental components. We investigated whether genetic susceptibility to T1D could be mediated by changes in DNA methylation, an epigenetic mechanism that potentially plays a role in autoimmune diabetes. Using data from a non-diabetic population comprising blood samples taken at birth (n=844), childhood (n=911) and adolescence (n=907), we evaluated the associations between 65 top GWAS single nucleotide polymorphisms (SNPs) and genome-wide DNA methylation levels. We identified 159 proximal SNP-cytosine phosphate guanine (CpG) pairs (cis), and 7 distal SNP-CpG associations (trans) at birth, childhood, and adolescence. There was also a systematic enrichment for methylation related SNPs to be associated with T1D across the genome, after controlling for genomic characteristics of the SNPs, implying that methylation could either be on the causal pathway to T1D or a non-causal biomarker. Combining the principles of Mendelian Randomization and genetic colocalization analysis, we provided evidence that at 5 loci, *ITGB3BP*, *AFF3*, *PTPN2*, *CTSH* and *CTLA4*, DNA methylation is potentially on the causal pathway to T1D.

## Introduction

Type 1 diabetes (T1D) is a polygenic disease with more than 50% of genetic susceptibility attributable to the human leukocyte antigen (HLA) class II region and the remaining attributable to the non-HLA region (1). Over the last decade, genome-wide association studies identified 62 independent loci and over 100 GWAS SNPs associated with T1D risk (2), but the biological pathways for most are unknown.

DNA methylation is an epigenetic event that occurs at cytosine – phosphate – guanine (CpG) residues and can be modified by both genetic and environmental exposures. Genetic and epigenetic interactions have been postulated to contribute to susceptibility to a number of autoimmune disorders (3; 4).

Previous work focused on identifying methylation differences among T1D monozygotic twins (5), allowing changes of DNA methylation levels that are introduced by environment to be captured (6-8), but no study has systematically investigated the causal relationship between genetically driven DNA methylation changes and T1D. Genetic variance has been reported to explains 24% of the methylation variance in childhood and 21% in middle age, when T1D can develop (9; 10). We therefore asked whether genetic susceptibility for T1D could be mediated by DNA methylation, which subsequently lead to altered gene expression and immune cell function. To address this question, the associations between T1D susceptible single nucleotide polymorphisms (SNPs) and DNA methylation need to be established first. When a T1D-SNP is associated with DNA methylation, four potential scenarios can occur:

(1) the SNP has a causal effect on T1D mediated by the changes of DNA methylation levels (Figure 1a);
(2) the SNP has a causal effect on T1D (i.e. via altering gene expression), which in turn alters DNA methylation levels (reverse causal, Figure 1b);
(3) the SNP that causes methylation change is in linkage disequilibrium (LD) with the causal variant of T1D (Figure 1c);
(4) the SNP influences methylation and T1D via separate mechanisms, an effect known as horizontal pleiotropy (Figure 1d).

**Figure 1.**
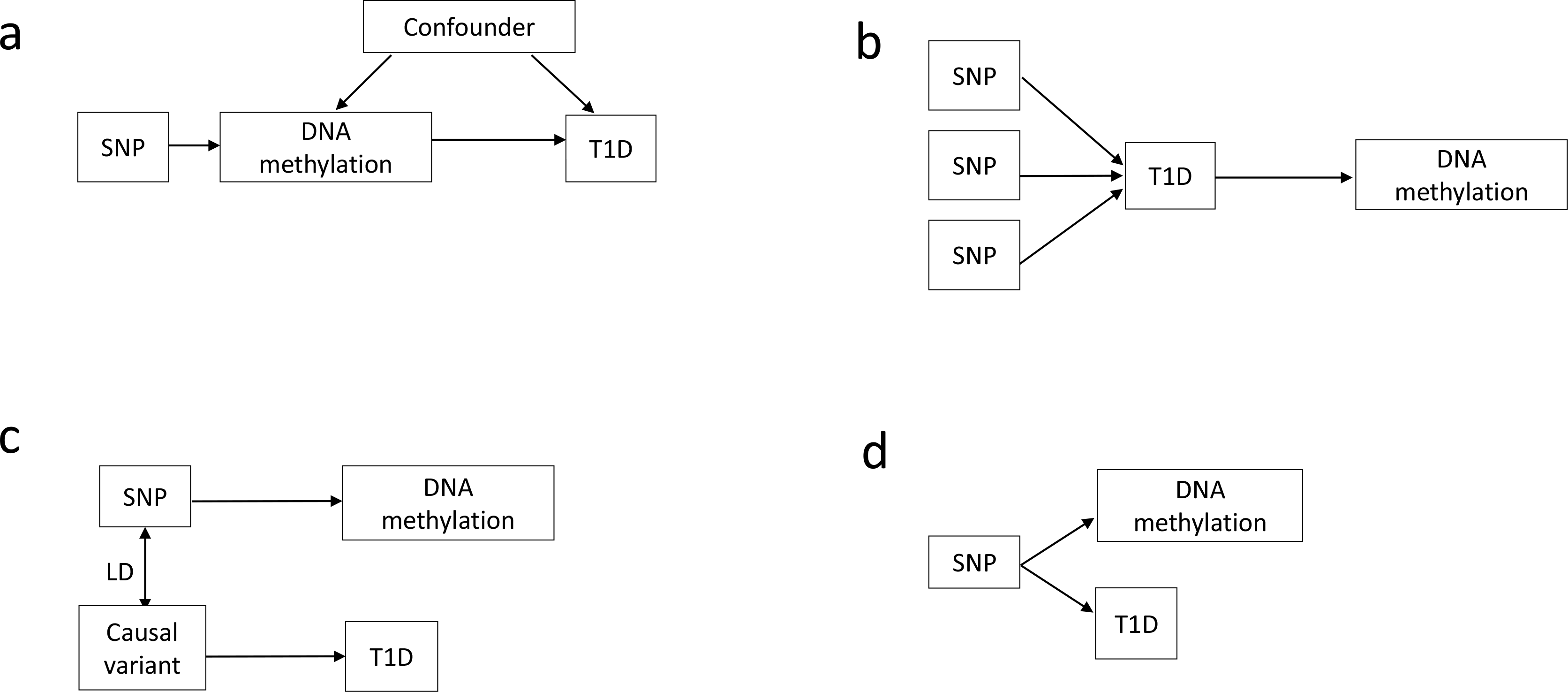
Four possible scenarios that could explain the associations between SNP, DNA methylation and T1D. a, DNA methylation mediates the genetic risk of T1D; b, SNPs first increases T1D liability, which in turn changes DNA methylation levels; c, a SNP that regulates DNA methylation could simply be in LD with another causal variant that influences T1D; d, a SNP is associated with DNA methylation and T1D via independent biological pathways (horizontal pleiotropy).

To disentangle the causal associations between DNA methylation and T1D, we employed principles of Mendelian Randomization (MR). MR is a statistical framework to infer causal relationships in hypothesis testing, using genetic polymorphisms to create a pseudo-experimental design. Since genotypes are randomly assigned at conception and not influenced by environmental confounders, they can be used as instruments to proxy exposures that are potentially influencing a trait, thereby mimicking a randomised controlled trial (11). In the context of DNA methylation, SNPs that regulate methylation levels at nearby CpG sites (defined within 1Mb distance, known as cis-mQTLs) can be used to investigate the causal effect of DNA methylation on a trait (12). If a CpG site mediates genetic risk of T1D, the casual effect of this CpG site can be interpreted as the change in log odds for T1D per unit increase in the CpG methylation level due to its associated SNP. Compared with traditional MR, where the effects of genetic instruments on exposure and on associated traits are measured in the same population (hence one-sample MR), Two-Sample MR (2SMR) has been developed to enable causal inference using summary statistics from GWAS alone, circumventing the requirement that DNA methylation levels and T1D status are measured in the same sample, enabling much larger sample sizes (11).

Firstly, in this study, we established the association between top T1D GWAS SNPs and DNA methylation by performing an epigenome-wide association (EWAS) analysis in a young non-diabetic population, where associations are likely to be on the causal pathway to T1D. Secondly, we tested whether there is an overall association between cis-mQTLs and T1D. Subsequently, following the framework outlined in Richardson et al 2017 (13), we combined the principles of 2SMR with genetic colocalization analysis to assess the four scenarios outlined above. Finally, we tested whether the findings can be replicated in an independent cohort with T1D patients and their relatives. A flow chart summarising the analysis procedure is shown in Figure 2.

**Figure 2.**
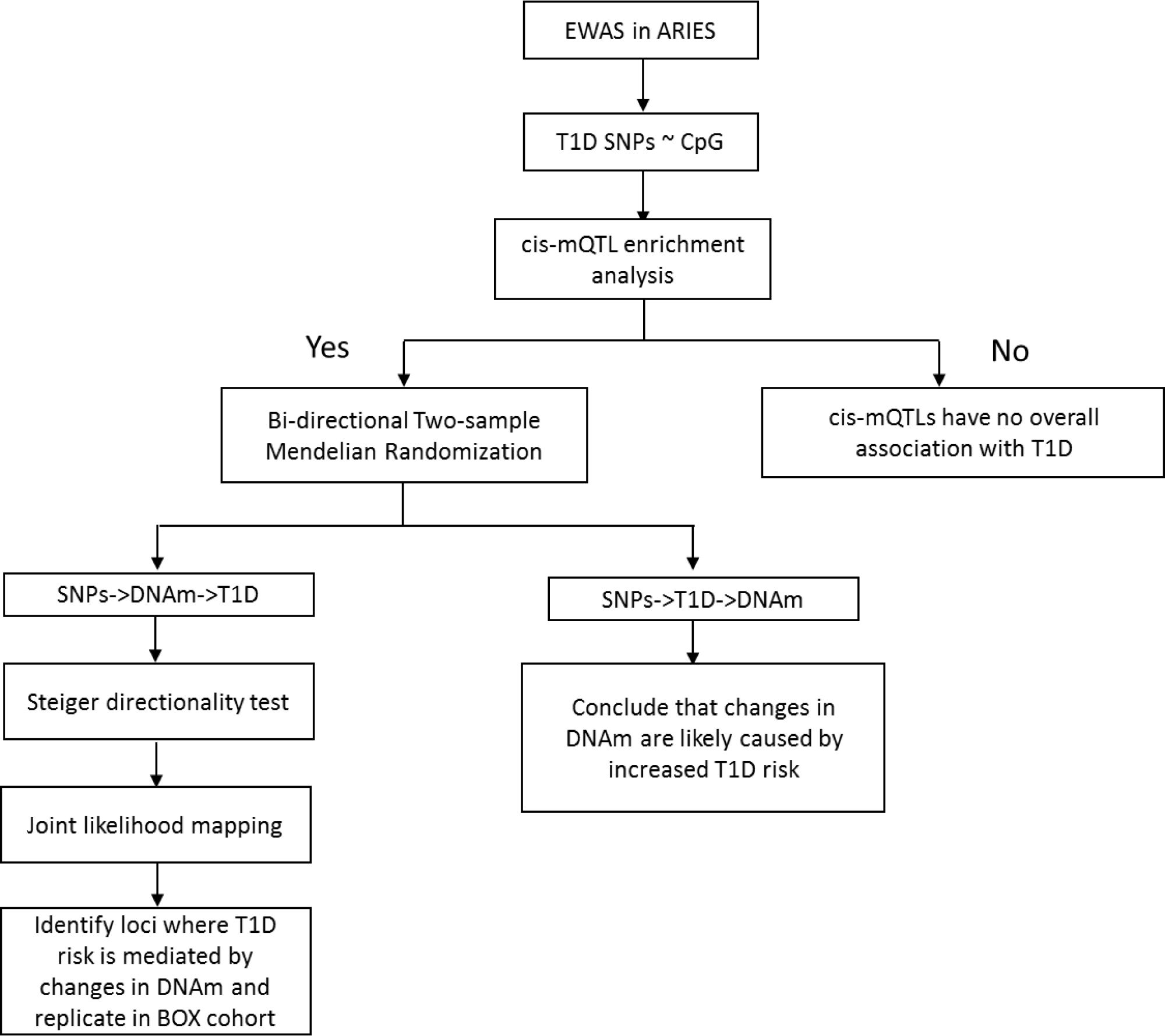
Flow chart summarising the overall analysis procedure in this study. EWAS: epigenome wide association analysis; DNAm: DNA methylation; mQTL: methylation quantitative trait loci

## Materials and methods

### DNA methylation data

#### Young non-diabetic population

DNA methylation data was obtained from the Avon Longitudinal Study of Parents and Children (ALSPAC) study, a large scale prospective study based in Avon, UK. ALSPAC recruited 14,541 pregnant women with expected delivery dates between 1^st^ April 1991 to 31^st^ December 1992, clinical data and biological samples were collected during pregnancy and at regular intervals postpartum from both parents and offspring (14; 15). The study website contains details of all the data that is available through a fully searchable data dictionary http://www.bris.ac.uk/alspac/researchers/data-access/data-dictionary/. DNA methylation data were generated either from whole blood or peripheral blood leukocyte samples, derived from 1,018 mother-offspring pairs using the Illumina HumanMethylation450 BeadChip (‘450k array’) (Illumina, San Diego, CA, USA) as part of the Accessible Resource for Integrated Epigenomic Studies (ARIES) project (16). The array quantifies DNA methylation levels of >485,000 CpG sites, which covers 99% RefSeq genes (17). The ARIES participants were selected based on the availability of DNA samples at two-time points for the mother (antenatal and at follow-up when the offspring were adolescents) and three-time points for the offspring (cord blood, childhood and adolescence). Methylation data from the offspring at adolescence (mean age 17.1 years, n=907), childhood (mean age 7.5 years, n=911) and birth (mean range, n=844) were used in this study. Detailed quality control and normalization procedures have been described previously (16). The final methylation data contained 459,734 probes per individual.

#### T1Dpopulation

As part of the methylation study, 16 families including 45 individuals were selected from the Bart’s Oxford family study of type 1 diabetes (BOX) (18). DNA from probands, their parents and grandparents were analysed using the 450k array, their clinical characteristics were summarized in Supplementary Table 1. In brief, DNA methylation was normalized using SWAN normalization and further corrected for batch effect, age, gender and cell heterogeneity using the Minfi and SVA package under the R statistical software (version 3.2.2), prior to plotting against genotype.

### Individual level genotype data

Individual level genotype data on the ARIES cohort were generated using Illumina HumanHap550-quad chips by Sample Logistics and Genotyping Facilities at the Wellcome Trust Sanger Institute and LabCorp (Laboratory Corporation of America). Details for quality control have been described before (19). SNPs (n=113) spanning 57 genomic regions that are associated with T1D at genome-wide significant level were obtained from immunobase.org, including six additional SNPs associated with T1D from a recent GWAS study (2). Among theses, sixty-seven available SNPs (LD r^2^ < 0.1) were selected and where necessary, proxy SNPs (minimal r^2^ = 0.6) were used to replace the original variants in order to obtain the required odds ratios for downstream MR analysis. Genotype data of sixty-five SNPs were available from ARIES, which were summarised in Supplementary Table 2.

Genotype data for the BOX participants were determined using Taqman^®^ allele discrimination assays (Life Technologies, UK). Taqman probes for rs2269242, rs9653442, rs3087243, rs3825932, and rs1893217 were purchased from Life Technologies (Thermo Fisher, UK).

### mQTL-CpG association analyses

To remove outliers, bimodally distributed beta values were converted to M values (20) and then rank-transformed into normal distributions. We regressed 65 independent SNPs with 459,734 CpG sites measured at adolescence, childhood and birth separately using a mixed effect linear model; age, gender, cellular compositions (21) were also included as covariates in the model. A Bonferroni threshold of 1.6×10^−9^ (0.05/65*465,877) was used to correct for multiple-testing. Analyses were performed using the MatrixEQTL package in R 3.2.2 statistical software on the University of Bristol High Performance Computing (HPC) cluster.

### Summary statistics

T1D GWAS summary statistics were obtained from two meta-analyses: one for the initial analysis (Data 1) (2) and one for replication (Data 2) (22); details were summarized in Supplementary Table 3. Data were retrieved from Immunobase.org; only *p*-values for Data 2 were available.

mQTL summary data was obtained from ARIES participants from a previous study (9), in which *p*-values of SNP-CpG associations were retrieved using the TwoSampleMR R package.

### mQTL enrichment analyses

We investigated whether there is an overall association between cis-mQTLs and T1D more than expected by chance. cis-mQTLs (LD r^2^<0.1) that have shown strong associations (p<1×10^−14^, Type I error rate 0.2%) with DNA methylation were retrieved from the ARIES cohort (9); their *p*-values for T1D associations were retrieved from the two GWAS summary statistics (Data 1 and Data 2, respectively). The enrichment of cis-mQTL could be either due to their (1) distinct SNP properties (allele frequency, LD, gene distance) or (2) due to distinct genomic locations (i.e. promoter, intron, exon, 5’ UTR, or 3’ UTR). To control for these two factors, null SNPs were selected to match cis-mQTLs in two ways. Firstly, null SNPs were chosen based on similarities in minor allele frequency (MAF) and LD structures (23). For example, null SNPs must be at least 1000kb away from mQTLs; the maximum MAF deviation of null SNPs from mQTLs is 0.02; and LD scores (24) of null SNPs are in the same quintile bin of mQTLs. Secondly, null SNPs were chosen based on similarities in genomic annotations, such as, intron, exon, 5’UTR, 3’UTR or promoter SNPs. For both methods, null SNPs were sampled without replacement. Fisher’s combined probability test was used to obtain an overall association with T1D:

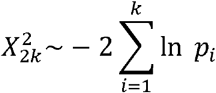

To generate a distribution for null SNPs, the same number of null SNPs was randomly drawn 10,000 times to obtain 10,000 combined *p*-values. The empirical *p*-value, reflecting the likelihood of observing a combined *p*-value at least as extreme as the combined *p*-value for mQTLs in the null distribution, is calculated by ranking all the 10,000 *p*-values for null SNPs (9).

### Bi-directional Two-Sample Mendelian Randomization

Forward 2SMR was used to test whether a CpG is causally influencing T1D (Figure 1a). To be considered as a valid instrument to proxy DNA methylation, first, a SNP must be strongly associated with DNA methylation; second, a SNP must only influence T1D via DNA methylation; third, a SNP must be independent of confounders of the methylation-T1D associations (i.e. hyperglycemia and medications) (25). To reduce potential pleotropic effect, only cis-mQTLs were used as instruments since trans-mQTLs may influence methylation via multiple biological pathways. To exclude potential instrument-confounder associations, we also ensured that instruments were not associated with fasting glucose concentration in a large GWAS meta-analysis involving 133,010 non-diabetic European individuals (26). The causal effect of CpG to T1D was then determined using the Wald ratio estimator, derived by the ratio between the log odds of SNP on T1D association and the coefficient of SNP on DNA methylation association (27).

To test the causal effect of T1D on DNA methylation, multiple instruments were used for individual CpG as an outcome (Figure 1b). For all CpGs, mQTL used as an instrument in the forward 2SMR were excluded. The causal effects of multiple SNPs were combined in a fixed-effect meta-analysis using MR – inverse variance weighting. To account for multiple testing in the outcome (i.e. CpGs are co-methylated if they are located close to each other on a chromosome), we used matrix spectral decomposition (matSpDlite) (28) to determine the number of independent tests in the outcome, it also reports an alpha level that is required to keep the Type I Error rate at 5% (28).

### MR-Steiger directionally test

In some situations, such as DNA methylation, it is difficult to ascertain whether genetic risk first causes changes in DNA methylation which subsequently results in T1D risk or vice versa. As a verification of 2SMR results, we used MR-Steiger to assess whether DNA methylation is the likely exposure and T1D risk is the likely outcome. MR-Steiger estimates the proportion of variance in the exposure and in the outcome that is explained by genetic instruments. Causal direction is then determined based on whether exposure variance or outcome variance is subject to the primary effect of SNPs (29).

### Bivariate fine-mapping

To estimate the probability of shared causal variants for DNA methylation and T1D (scenario 3, Figure 1c), we performed joint likelihood mapping (JLIM) (30). Concordance between top SNPs for the two traits would suggest that DNA methylation potentially reside in the causal pathway to T1D risk. The concordance rates were determined after accounting for chance, under 1000 permutations.

## Results

### The association between DNA methylation and T1D susceptible SNPs across three-time points of life

Of the 65 independent T1D GWAS variants, thirty-eight SNPs were consistently found to associate with a total of 166 CpG sites under the Bonferroni corrected threshold *(p* < 1.6e-9) at adolescence, childhood and birth in the ARIES cohort. Seven SNP-CpG pairs were in trans (distance >1Mb), the remainder in cis; these data are summarised in Supplementary Table4. Figure 3 shows the genomic distribution of 166 total CpGs, including extensive associations at the HLA locus (Supplementary Figure1). According to 450k array annotation, approximately 45% CpGs were located in the gene body / introns; 24% in promoters / distal promoters. Methylation variance (R^2^) explained by T1D SNPs varied largely, the strongest association lies between rs7149271 and cg20045882 on chromosome 14, where rs7149271 explained greater than 78% methylation variance across all three-time points. At the HLA locus, rs3104163 is in high LD with rs9272346 (r^2^=0.84), a variant most strongly associated with T1D but not available in our summary statistics (OR=18.5). rs3104363 regulates 70 CpGs within the HLA locus and the most strongly associated CpG site is cg01889448, for which rs3104363 explained at least 53% methylation variance across three-time points. The overall effect sizes of T1D variants on DNA methylation levels are consistent across three-time points, where the correlation between adolescence and childhood is 0.996 (95% CI: 0.995, 0.997, *p* < 2.2e-16); between adolescence and birth is 0.984 (95% CI: 0.979, 0.989, *p* < 2.2e-16); and between childhood and birth is 0.987 (95% CI: 0.982, 0.990, *p* < 2.2e-16).

**Figure 3.**
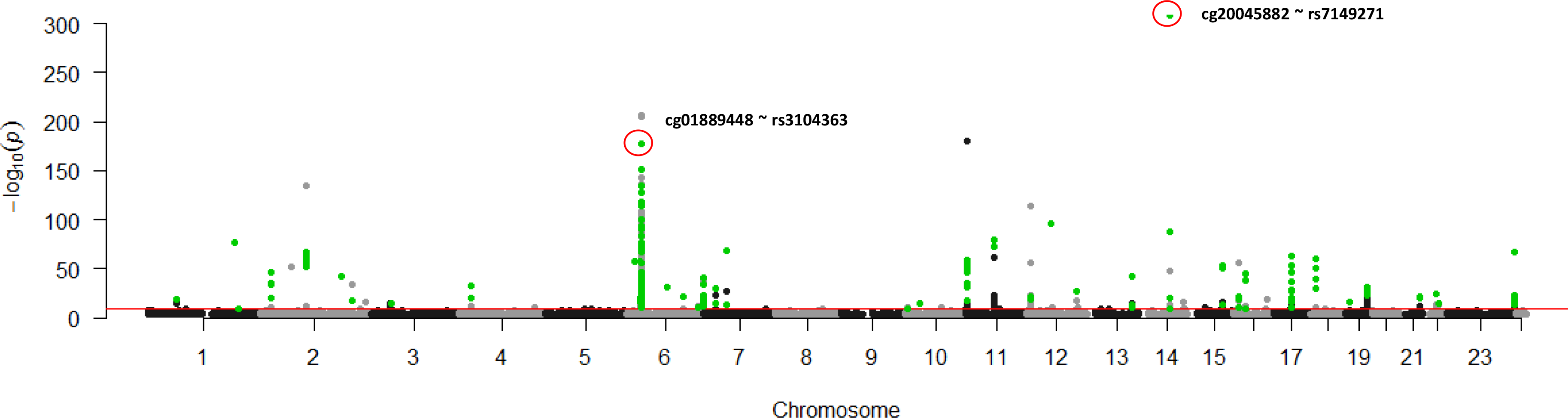
Genomic distribution of CpG sites that are associated with T1D GWAS variants. Manhattan plot showing the CpG sites associated with 38 T1D GWAS variants above the Bonferroni threshold 1.6e-9 (redline); green highlighted dots were those (n=166) that were consistently detected at adolescence, childhood and birth; there is a peak reflecting mQTL-CpG association at the HLA locus (chromosome 6).

### Genome-wide enrichment of cis-mQTLs and Type 1 diabetes associations

As a primary analysis, we overlapped adolescent cis-mQTLs with an initial discovery dataset (Data 1). These cis-mQTLs were evenly distributed across the genome (data not shown) and were devoid of HLA-SNPs (chr6: 28,477, 797-33, 448,354, hg19). Their overall associations with T1D were significantly enriched in SNPs with low GWAS *p*-values, matching null SNPs to cis-mQTLs either by SNP properties (Figure 4 a) or by genomic annotations (Figure 4 b). Secondary analyses using childhood and neonatal cis-mQTLs revealed the same findings (Supplementary Figure2). To verify these observations, we performed a replication study using Data 2. Compared to Data 1, there was a stronger enrichment when cis-mQTLs were matched either by SNP properties or by genomic annotations (Supplementary Figure2). These data suggest that there is a shared genetic influence of DNA methylation levels and T1D. Furthermore, from each of the enrichment analyses we identified a number of cis-mQTLs that have smaller observed GWAS p-values than theoretical p-values (the probability of T1D association by chance). Most of these cis-mQTLs lie within known T1D susceptible loci and are in strong to moderate LD with index SNPs. However, rs605093 resides at Chr11q24.3 that was not reported by previous GWA studies or in LD with any index variants (*p* _observed.meta_ = ^4.22e-6^, *p* _theoretical_ = 1.01e-5).

**Figure 4.**
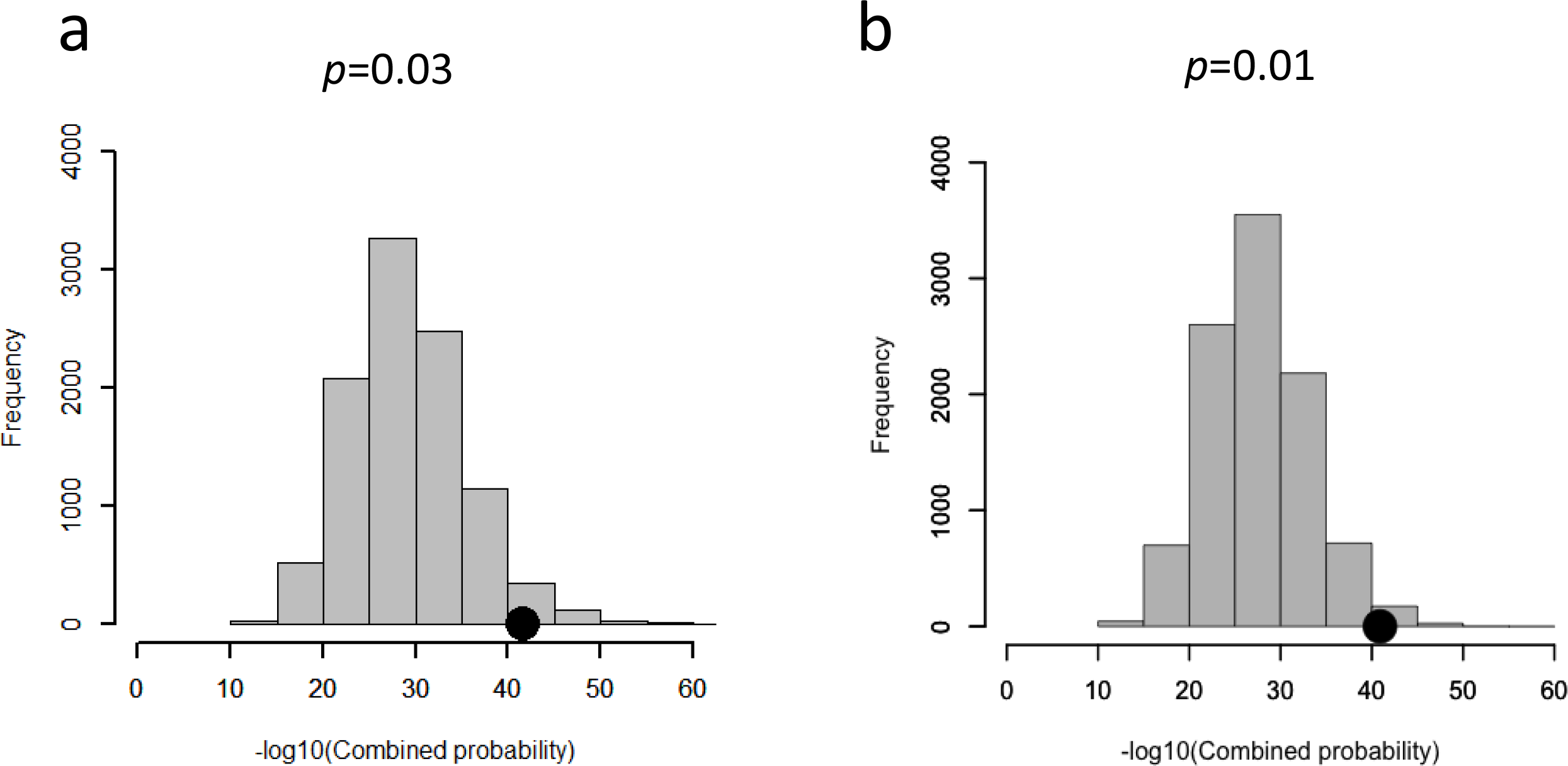
cis-mQTLs are enriched in SNPs with low GWAS *p*-values associated with T1D. a, A representative plot showing the enrichment analysis conducted using the adolescent data, when null SNPs were matched to cis-mQTLs via SNP properties; b, when null SNPs were matched to cis-mQTLs via genomic annotations. T1D GWAS p-values were extracted from meta-analysis Data 1.

### DNA methylation potentially mediates T1D genetic risk

After removing SNPs with harmonization issues (31), we obtained one hundred and twenty-eight CpG associations unique to 33 cis-mQTLs for analyses. Forward 2SMR showed that the Wald ratios for all the 128 CpGs were significant and the effects were consistent at adolescence, childhood and birth (*p* < 0.05, Supplementary Table5).

In the reverse 2SMR, ‘matSpDlite’ estimated 124 independent CpGs in the outcome and a p-value of 4.12e-4 is required to keep the type I error rate at 5%. At this threshold, there was no evidence for reverse causation at all three-time points. It is however important to note that the statistical power to detect an effect in this direction is low because the outcome sample size was small. These data are summarised in Supplementary Table5.

### MR-Steiger test to verify the direction of causality

MR-Steiger at the forward direction showed that for most CpGs within the HLA region, SNPs explained methylation variance more than T1D variance (36 out of the 64 CpGs indicated methylation as the exposure consistent at three-time points, *p* < 0.01, sensitivity ratio >2.35, Supplementary Table6). Outside the HLA, the same causal direction was inferred for all the CpGs and this effect was consistent at all three-time points (Supplementary Table6). We did not test the reverse direction due to insufficient statistical power to detect an effect.

### Bivariate fine-mapping pinpointed overlapping methylation and T1D causal variants

In the 32 non-HLA loci, JLIM analyses pinpointed shared causal variants in 5 loci, mediated by 10 CpG sites (*p* < 1e-3, Table 1, Figure 5). CpGs in the HLA region were however excluded from the analysis owing to the extensive LD structure and high false discovery rate (30).

**Table 1.**
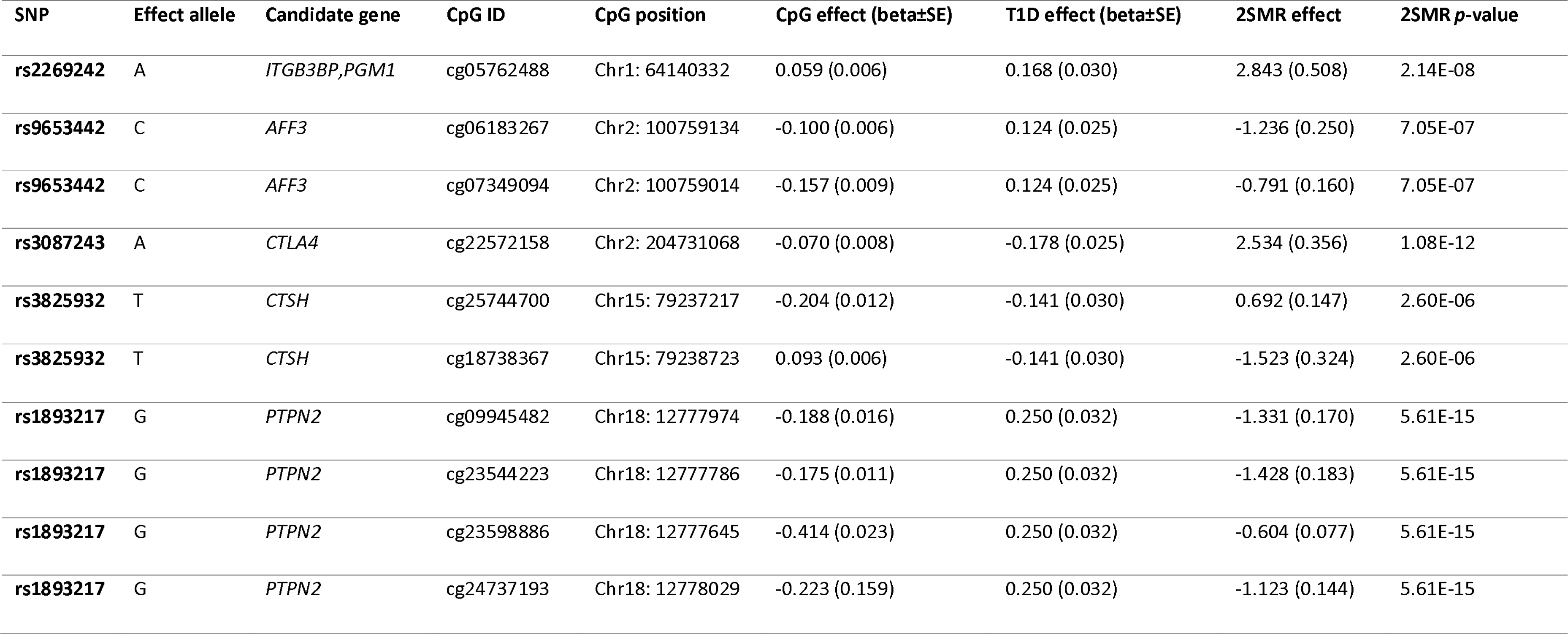
Representative 2SMR results from the adolescence dataset that survived the Joint Likelihood Mapping. CpG effect denotes the addition of effect allele relative to other allele on CpG methylation changes (beta coefficient ±SE); T1D effect denotes the addition of effect allele relative to other allele on T1D risk (beta coefficient ±SE, beta coefficient equals log odds ratio); 2SMR effect denotes the change of log odds on T1D per unit increase in DNA methylation due to its associated SNP.

| SNP | Effect allele | Candidate gene | CpG ID | CpG position | CpG effect (beta±SE) | T1D effect (beta±SE) | 2SMR effect | 2SMR p-value |
| --- | --- | --- | --- | --- | --- | --- | --- | --- |
| rs2269242 | A | *ITGB3BP, PGM1* | cg05762488 | Chr1: 64140332 | 0.059 (0.006) | 0.168 (0.030) | 2.843 (0.508) | 2.14E-08 |
| rs9653442 | C | *AFF3* | cg06183267 | Chr2: 100759134 | -0.100 (0.006) | 0.124 (0.025) | -1.236 (0.250) | 7.05E-07 |
| rs9653442 | C | *AFF3* | cg07349094 | Chr2: 100759014 | -0.157 (0.009) | 0.124 (0.025) | -0.791 (0.160) | 7.05E-07 |
| rs3087243 | A | *CTLA4* | cg22572158 | Chr2: 204731068 | -0.070 (0.008) | -0.178 (0.025) | 2.534 (0.356) | 1.08E-12 |
| rs3825932 | T | *CTSH* | cg25744700 | Chr15: 79237217 | -0.204 (0.012) | -0.141 (0.030) | 0.692 (0.147) | 2.60E-06 |
| rs3825932 | T | *CTSH* | cg18738367 | Chr15: 79238723 | 0.093 (0.006) | -0.141 (0.030) | -1.523 (0.324) | 2.60E-06 |
| rs1893217 | G | *PTPN2* | cg09945482 | Chr18: 12777974 | -0.188 (0.016) | 0.250 (0.032) | -1.331 (0.170) | 5.61E-15 |
| rs1893217 | G | *PTPN2* | cg23544223 | Chr18: 12777786 | -0.175 (0.011) | 0.250 (0.032) | -1.428 (0.183) | 5.61E-15 |
| rs1893217 | G | *PTPN2* | cg23598886 | Chr18: 12777645 | -0.414 (0.023) | 0.250 (0.032) | -0.604 (0.077) | 5.61E-15 |
| rs1893217 | G | *PTPN2* | cg24737193 | Chr18: 12778029 | -0.223 (0.159) | 0.250 (0.032) | -1.123 (0.144) | 5.61E-15 |

**Figure 5.**
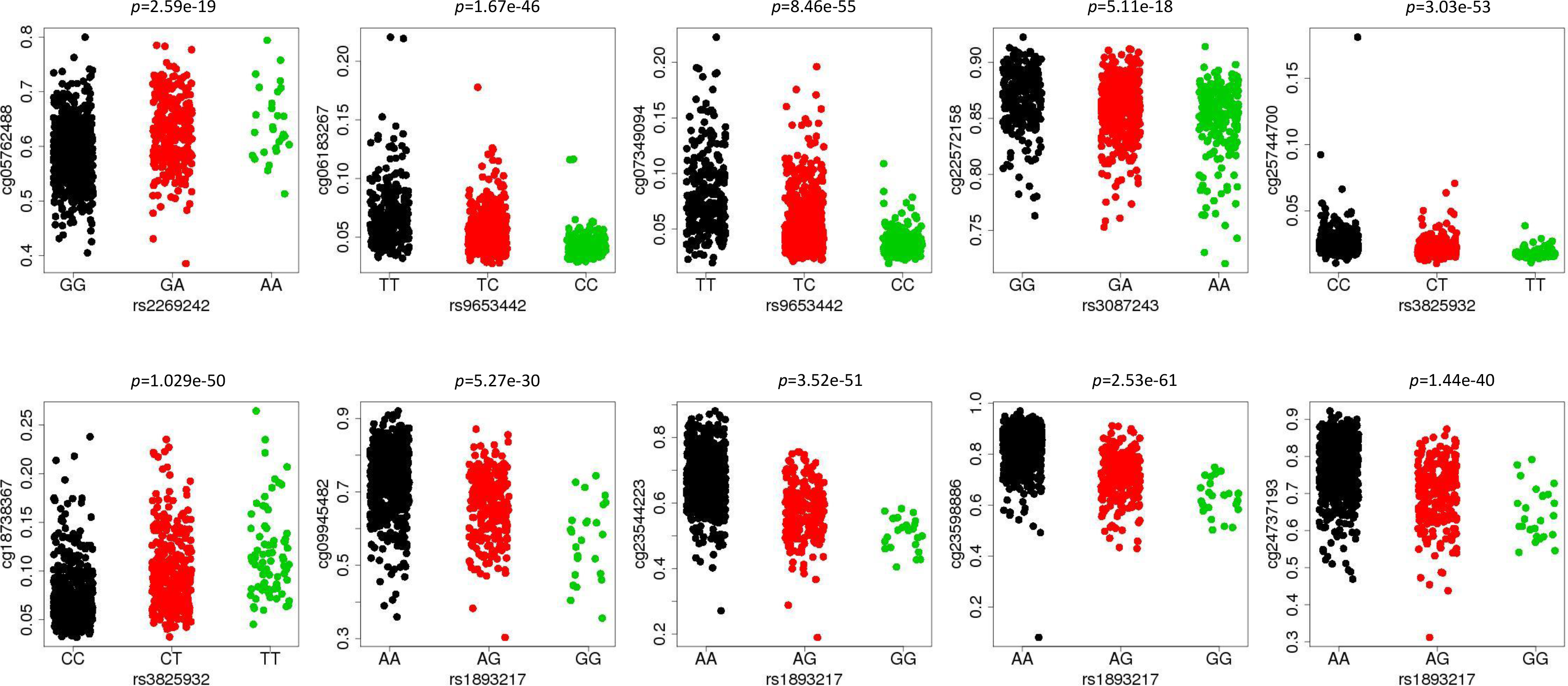
DNA methylation levels of CpG sites and their associations with T1D GWAS variants that survived JLIM analyses, obtained from the ARIES adolescent participants. Y - axis represents beta values for each CpG site. The inner most genotype in X - axis is comprised of two other alleles, the outer most genotype is comprised of two effect alleles.

### Replication in T1D cohort

The 10 SNP - CpG associations were further assessed in 45 individuals participating the BOX study, where methylation 450k array data were available. Nine out of ten associations showed similar patterns comparing to the ARIES cohort, after fitting the DNA methylation levels and SNP genotype into linear regression models (Figure 6). However, given the small sample size of the replication cohort, there is insufficient power to obtain significant *p*-values for cg05762488 and for cg025744700 (Figure 6).

**Figure 6.**
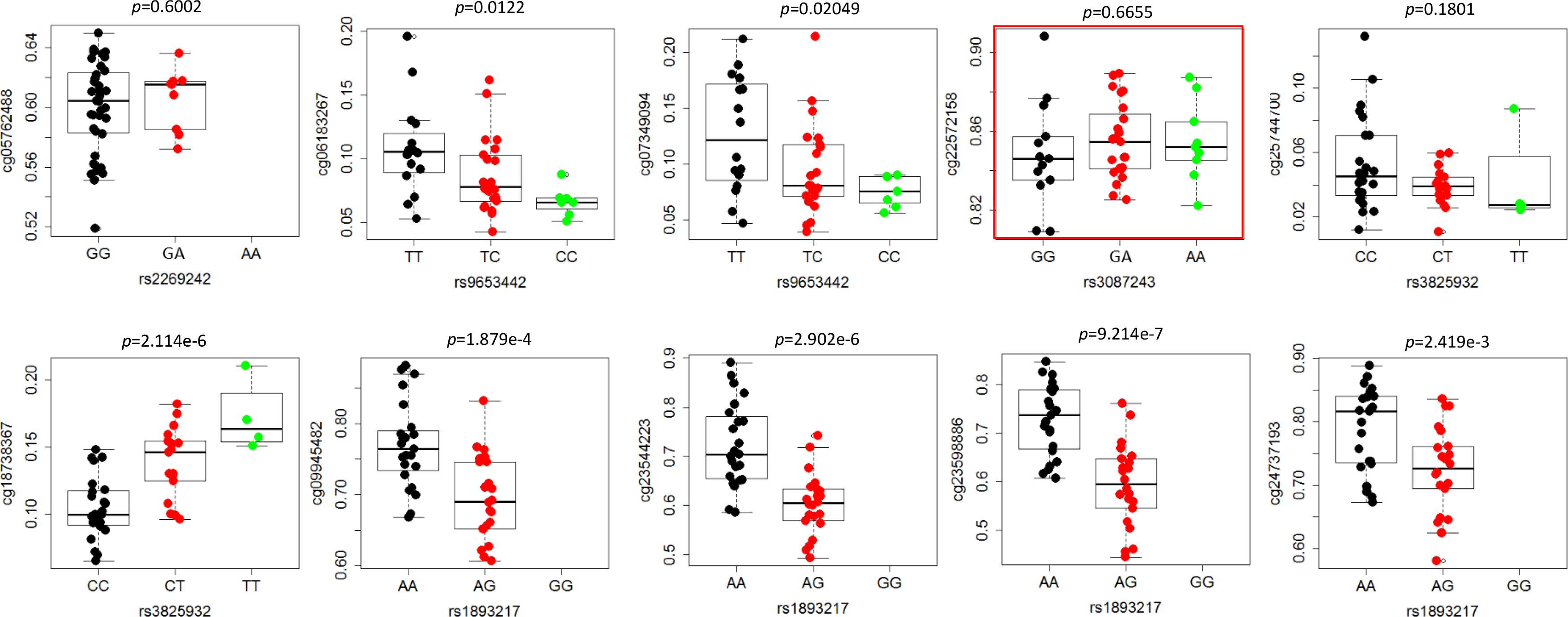
Replication of SNP and DNA methylation associations in the Bart’s oxford T1D cohort. 45 individuals were analysed in the BOX cohort. Nine out of ten SNP-CpG pairs showed similar associations compared to the ARIES participants. The SNP – CpG pair that did not replicate the ARIES result was highlighted in red.

## Discussion

One hypothesis of the mechanisms underlying T1D is that genetic variants alter DNA methylation levels, which in turn influence genes that are essential to immune tolerance as well as beta cell function, increasing the risk of T1D. To the best of our knowledge, this is the first study that systematically evaluated causal associations between methylation and T1D in a large-scale population.

Our EWAS identified widespread genetic and epigenetic interactions in known T1D susceptible loci; more than half of the T1D-SNPs were associated with proximal methylation variation in the genome, of which 40% resides at the HLA locus. More importantly, we showed genome-wide enrichment of cis-mQTLs and T1D associations in a young non-diabetic population at three-time points of life, implying that theses intrinsic methylation heterogeneities could either be causally influencing the liability of T1D or facilitate as biomarkers for islet autoimmunity development and progression.

A previous analysis based on SNP heritability however did not show significant association between cis-mQTLs and T1D (9). Narrow sense heritability of T1D was estimated to be approximately 0.8 (32; 33). The lack of enrichment of cis-mQTLs in SNP heritability found by the previous study, was probably due to limited number of cis-mQTLs used in the estimation. Since approximately 20,000 cis-mQTLs were identified in the ARIES study (9), more cis-mQTLs are perhaps required to capture enough genotypic variance to explain a highly heritable condition. We found that rs605093 was associated with T1D more than expected by chance. rs605093 was detected as one of the most associated T1D SNPs but failed to reach genome-wide significance, although no follow-up study was performed to confirm its true association (34). Since GWAS tend to omit variants with small effect sizes due to the burden of multiple testing, our data may suggest a weak effect of rs605093 on T1D. This SNP is located at the intron 1 of *FLI-1* (Friend leukemia integration 1 transcription factor), which overlaps with a known susceptible region for rheumatoid arthritis (RA) and systemic lupus erythematosus (SLE).

Given that associations between methylation and T1D are subject to confounding and reverse causation, we applied the principles of Mendelian Randomization and JLIM to evaluate their causal relationships. Although at majority of the loci, DNA methylation appeared to be non-causal, our analysis pinpointed 5 loci, where T1D risk is potentially mediated by methylation variations at 10 CpG sites. Interrogating regulatory features from the ENCODE/Roadmap consortium revealed and enrichment of CpGs with chromosome regulatory elements (Supplementary Figure3). For example, the rs2269242 (tagging *ITGB3BP)* associated CpG (cg05762488) is located within a dense DNase I hypersensitive region and transcription factor binding site adjacent to gene *PGM1* (phosphoglucomutase −1) and upstream of the gene *ITGB3BP* (integrin beta 3 binding protein beta3-endonexin). *ITGB3BP* is a new candidate gene for T1D (2) and is involved in signalling pathways of apoptosis (35). The two CpGs (cg06183267 and cg07349094) associated with rs9653442 are situated in the exon1 of *AFF3* (AF4/FMR2 Family Member 3), which is a region enriched with DNase I hypersensitivity and H3K4me3 (associates with enhancers) signals in a range of immune cells. *AFF3* is a risk gene for rheumatoid arthritis (RA) (36), juvenile idiopathic arthritis (37) and T1D (2; 38) and may be involved in lymphoid development and plasma cell differentiation (39; 40). The two CpGs (cg25744700 and cg18738367) associated with rs3825932 are located in intron 1 and 5’ upstream of the gene *CTSH* (Cathepsin H), respectively. cg25744700 overlaps with a H3K27Ac peak (marks active chromatin), a DNase I cluster, as well as a transcription factor binding region; cg18738367 co-localizes with a DNase I cluster. Lowered gene expression of *CTSH* in beta cells has been correlated with increased beta cell apoptosis upon cytokine exposure as well as faster diabetes progression (41). The detailed functional effects of these methylation variations require follow-up laboratory investigations.

To support our data, we observed similar association in 9 out of 10 SNP-CpG pairs in the independent BOX cohort containing T1D probands and their relatives. Although the sample size of this cohort is small (n= 45) and there is a wide age range in the participants, the high replication rate in methylation patterns warrants our initial observations in ARIES.

There are several limitations of our study. First, given single genetic instruments used in the forward 2SMR, we were unable to robustly distinguish mediatory effect from horizontal pleiotropy (scenario 4, Figure 1d) without further support from experimental proof. Second, MR and its extension MR-Steiger test both suffer from insufficient statistical power to detect an effect in the reverse direction. This is because the sample size available for SNP effects on CpG levels was small (approximately 1000 individuals in the ARIES participants per time point). Third, although there is a strong evidence of shared causal variants for methylation and T1D at 5 loci, JLIM does not report which SNPs are the shared causal variants. Fourth, all our analyses were performed using whole blood or peripheral blood lymphocyte samples. As the mQTL effects are likely to be tissue and cell type specific (42), attempts to interpret our findings in a different tissue must be conducted with caution. Another limitation is that the 450k array typically covers 2% of the entire epigenome (17) and we were unable to investigate causal effects of uncovered CpG sites. It is thus possible that methylation influences T1D risk beyond the 5 loci identified in this study; those effects could not be tested without large-scale methylation sequencing data. We also reported mQTLs that regulate DNA methylation consistently at birth, childhood and adolescence, covering the spectrum of life when diagnosis of T1D peaks. It might be possible that some mediatory effects of DNA methylation are only specific to a particular time point (9). These changes are beyond the scope of the current analyses, future studies may focus on investigating how epigenetic regulations change during immune development.

In the HLA region, we observed extensive methylation associations with a top GWAS variant rs3104363 (T1D OR=3.7). A recent study reported over 180 CpGs in the HLA region that were differentially methylated dependent on the T1D susceptible DQ2/DQ8 haplotype (43). Our focus was on SNPs, but 17 haplotype dependent CpGs were also strongly associated with rs3104363 in our data. Of these 13 CpGs demonstrated evidence of mediating T1D risk. However, due to high false positive rates at the HLA locus we could not use JLIM to exclude LD confounding effects (30). Functional approaches are required to better understand the effects of SNP and haplotype dependent methylation on HLA expression.

In conclusion, the identification of putative genetically driven DNA methylation changes provides a rich source for follow-up verifications, as dissecting the functional effect of genetic and epigenetic interactions may help uncover novel mechanisms that contribute to the risk of T1D development.

## Acknowledgement

We are extremely grateful to all the families who took part in this study, the midwives for their help in recruiting them, and the whole ALSPAC team, which includes interviews, computer and laboratory technicians, clerical workers, research scientists, volunteers, managers, receptionists and nurses. We also sincerely appreciate the participants of the BOX study, nurses, administrators, and laboratory technicians for their contributions. The authors are grateful for Prof. John A Todd and Mr. Jamie Inshaw who kindly provided T1D GWAS summary statistics, as well as Prof. Yaron Tomer for his valuable advice on methylation data interpretation.

## Web Resources

Matrix eQTL http://www.bios.unc.edu/research/genomic_software/MatrixeQTL/;

2SMR and MR-Steiger https://github.com/MRCIEU/TwoSampleMR;

mQTL enrichment analysis https://github.com/olegkagan/Ye-et-al.-2017;

JLIM https://github.com/cotsapaslab/jlim;

matSpDlite

https://github.com/snewhouse/BRC_MH_Bioinformatics/blob/master/misc_sh/matSpDlite.R;

Immunobase http://www.immunobase.org;

UCSC genome browser https://genome.ucsc.edu/

## Research funding

The UK medical Research Council and Wellcome (Grant ref: 102215/2/13/2) and the University of Bristol provide core support for ALSPAC. GWAS data were generated by Sample Logistics and Genotyping Facilities at the Wellcome Trust Sanger Institute and LabCorp (Laboratory Corporation of America) with support from 23andMe. Methylation data in the ALSPAC cohort were generated as part of the UK BBSRC-funded (BB/I025751/1 and BB/I025263/1) Accessible Resource for Integrated Epigenomic Studies (ARIES). Methylation data used in the Bart’s Oxford Study (BOX) was generated using funding from Diabetes UK (14/0004869). This publication is the work of the authors; and J.Y will serve as guarantor for the contents of this paper. J.Y. was funded by a Diabetes Wellness & Research Foundation non-clinical fellowship N-C/2016/Ye. T.G.R. was supported by the Elizabeth Blackwell Institute Proximity to Discovery award EBI 424. M.S. was supported by the Economics and social research council ES/N000498/1. G.H. was supported by the Medical Research Council MC_UU_12013/1-9.

## Abbreviations

ARIES: Accessible Resource for Integrated Epigenomic Studies
ALSPAC: Avon Longitudinal Study of Parents and Children
BOX: Bart’s Oxford family study of Type 1 Diabetes
CpG: Cytosine-phosphate-guanine dinucleotides
GWAS: Genome-wide association study
HLA: Human leukocyte antigen
JLIM: Joint likelihood mapping
LD: Linkage disequilibrium
SNP: Single nucleotide polymorphism
T1D: Type 1 diabetes
MAF: Minor allele frequency
MR: Mendelian Randomization
mQTL: methylation quantitative trait loci
RA: Rheumatoid arthritis
2SMR: Two Sample Mendelian Randomization

